# Functional composition of subsoil microbial communities changes with oak mortality

**DOI:** 10.1101/2024.11.28.625745

**Authors:** Anna C. Goodman, Ernest N. Walker, Glade D. Bogar, Laura M. Bogar

## Abstract

Tree mortality in oak savannas is increasing under climate change, but its impact on microbial communities and soil carbon below the top 20 centimeters is relatively unknown. Deep tree roots, their ectomycorrhizal fungi, and associated bacteria may have a particularly important effect on landscape carbon storage, as they mediate the transfer of recently fixed plant carbon into deep soil and subsoil layers. To investigate how tree mortality impacts microbes and soil carbon, we sampled under living and recently dead *Quercus douglasii* trees in a California oak savanna, gathering depth-resolved soil cores to 45 cm below the surface. We captured finely resolved biological detail on these soil samples, comparing living (RNA-based) to potential and historical (DNA-based) microbial communities and assessing microbial biomass with phospholipid fatty acid analysis. Tree mortality greatly reduced the abundance of ectomycorrhizal fungi, particularly in subsoils. Fungal niches were more variable at depth under dead trees than under living ones, and RNA-based profiling captured substantially different communities than DNA, especially under living trees. However, tree mortality three years prior to our study did not impact the overall quantity of carbon stored in the soil. Tree mortality can have profound effects on the interactions between tree roots, mycorrhizal fungi, and soil bacteria, which may shift soil carbon dynamics over long time scales. Understanding the mechanisms of these interactions, and their time scales, will improve our ability to predict and manage soil carbon in savanna landscapes as drought and heat events kill more oaks in arid climates.

**Highlights:**

- Ribosomal RNA, from living cells, revealed different communities than from DNA.
- Microbial functions changed more with depth and tree health than taxonomy.
- Differences were most extreme below 20 cm depth.
- Microbial population distributions changed under living and dead trees.
- Total microbial biomass and carbon were similar beneath living and dead trees.

## Introduction

In California, increasing drought and fire severity are prompting huge waves of tree mortality. Tree cover loss has been as high as 10% per major drought event in blue oak (*Q. douglasii*) (Huesca et al., 2021), a dominant species in much of California’s oak savanna. Soils in these savannas contribute notably to global soil carbon storage, sequestering an estimated 110 g organic C/m^2/year (Ma et al., 2016). This soil organic carbon (SOC) consists largely of plant and microbial biomass, the persistence of which is determined by the balance of inputs (the deposition of plant-fixed atmospheric carbon into the soil) and outputs (respiration by roots and soil biota, often during microbial decomposition of existing soil organic matter) (Davidson and Janssens, 2006). Despite their role in carbon release, increased abundance of microbes may improve overall soil carbon storage capacity, as microbial carbon is more persistent and less vulnerable to degradation than plant carbon (Cotrufo et al., 2019). While SOC input/output dynamics are the subject of considerable research, existing research largely ignores SOC stored at depths of 30cm or below (Richter and Billings, 2015) where emerging studies assert that biogeochemical controls on C cycling are distinct from those at the surface (Dove et al., 2021). At these depths, deep roots are likely to control SOC dynamics; live roots deposit recently-fixed carbon and promote larger microbial populations in deep soils (Gocke et al., 2017). Tree roots may be particularly influential in deep soils, as their positive effect on the population size of soil microbiomes has been shown to be even greater at depth than in surface soils (Beule et al., 2022). Californian oak savannas have been observed to store up to 50% of soil carbon below 30cm (Moreland et al., 2021), so the activity of dominant trees such as *Q. douglasii* is likely to play a crucial role in deep soil carbon storage. Little is known about how the death of these trees will alter SOC storage dynamics. In coniferous forests, local tree mortality has been associated with a rapid and dramatic release of soil-stored carbon and tractable shifts in soil microbial community composition (Xiong et al., 2011), while by contrast, below evergreen oaks, defoliation and mortality is associated with increased SOC (García-Angulo et al., 2020). None of these studies have sampled deep soils, however, so the strongest effects of tree mortality may have been overlooked.

In many tree species, the influence of deep roots on SOC may be mediated by ectomycorrhizal fungi, which dominate fungal communities near tree hosts throughout the A, B, and C horizons (Dove et al., 2021; Beule et al., 2022). Ectomycorrhizal fungi form close mutualisms with the roots of many plants – including oaks – through which they can assimilate recently fixed, plant-derived C while mineralizing soil-bound nutrients and delivering them to plant hosts (Becquer et al., 2019; Zhou et al., 2022). The subsequent translocation of plant-derived C throughout ectomycorrhizal hyphae may extend trees’ metabolic sphere of influence beyond the rhizosphere into the more spatially distributed “hyphosphere” (Zhou et al., 2022). There, the presence of ectomycorrhizal fungi may improve SOC stability: the “Gadgil effect” posits that ectomycorrhizal fungi limit soil organic matter (SOM) decomposition by competing with saprotrophic fungi for limited N resources (Gadgil and Gadgil, 1975; Fernandez and Kennedy, 2016), and tree association with ectomycorrhizal fungi has been correlated globally with high soil C:N ratios (Averill et al., 2014; Cotrufo et al., 2019).

Ectomycorrhizal fungi may also support SOC stability by facilitating the efficient transduction of young SOC into stable mineral-associated organic matter (MAOM). Given that MAOM is predominantly composed of microbial biomass (Kögel-Knabner et al., 2008) and that local availability of plant-derived carbon is a primary driver of microbial proliferation in deep soils (Dove et al., 2021), the distribution of plant-derived carbon by ectomycorrhizal hyphae throughout the soil area could support large microbial communities in hyphosphere soils, resulting in large fractions of MAOM. A recent greenhouse bioassay affirms that ectomycorrhizal fungi can transform the taxonomic and functional composition of soil bacterial communities (Berrios et al., 2024) – which is of particular interest because bacterial biomass consistently dominates the MAOM fraction across ecosystems (Yu et al., 2022). *In vitro*, ectomycorrhizal fungi increase the alpha diversity of bacterial species and select for bacterial taxa with the genetic equipment to digest carbon-containing fungal metabolites (Berrios et al., 2024). This implies that exudates from ectomycorrhizal fungi may be an important food source for bacteria in soils where they are abundant, positioning ectomycorrhizal fungi as a key conduit for the drawdown of atmospheric carbon into stable bacterial biomass, and further implicating ectomycorrhizal host trees as key contributors to SOC input. However, the conditioning of bacterial communities by ectomycorrhizal fungi has seldom been documented in the field (Vik et al., 2013), and never in subsoils, so its prevalence and potential impact on real-world SOC storage cannot yet be extrapolated. Furthermore, extrapolation of microbial community response to tree mortality, depth, or ectomycorrhizal fungi may be hindered by the use of DNA metabarcoding, which fails to distinguish live, active microbes from the remnants of those which were present only historically (Carini et al., 2016).

The goal of this study was to illuminate the effects of deep-rooted, ectomycorrhizal tree mortality on soil carbon storage in a Californian temperate savanna. We aimed to investigate whether mortality in *Q. douglasii* shifts the size and composition of active subsoil microbial communities—specifically, with respect to mycorrhizal fungi and their bacterial associates—and whether these shifts correlate with a reduction in the carbon storage of related soils. To accomplish this, we compared both PLFA-based microbial abundance and DNA- and RNA-based microbial community composition between soils collected under living and fire-killed *Q. douglasii* across a depth gradient of 0-45cm. We hypothesized that living oaks and their associated mycorrhizae create niche space for a large, distinct microbial community across soil depths, which in turn increases soil carbon storage. We therefore predicted that soils associated with living trees would contain larger microbial populations than soils associated with dead trees, and that the size of these populations would remain more constant over depth. We further predicted that soils associated with living trees would contain larger populations of ectomycorrhizal fungi, and that observations of these fungi would be correlated with increased bacterial species richness and altered bacterial community composition. Finally, we predicted that soils harboring large populations of ectomycorrhizal-associated microbes would store more carbon.

## Methods

### 2.1 Site Description

The study was conducted at the University of California Natural Reserve System, Quail Ridge Reserve, California (38.478 N, 122.145 W) (Reserve DOI: 10.21973/N30T0K). Quail Ridge is a hilly peninsula with extreme topography, surrounded to the west, north, and south by the southern shores of Lake Berryessa. The ridge is in the Inner Coast Range, formed of lower Cretaceous through upper Jurassic subducted and uplifted marine sediments, comprising mostly fine-grained mudstone and shale, with rare conglomerates (Alt et al., 2016). Soils mapped in the sampling area include Bressa-Dibble Complex (30-50% and 50-75% slopes) and the Maymen-Millsholm-Lodo association (Lambert and Kashiwagi, 1978). Topography is the greatest determinant of soil development and plant community on the ridge, with a patchwork of oak woodland, savanna, grasslands, and chaparral demarcated by stark and sudden transitions in slope and aspect. In August 2020, the LNU Complex Fire burned the entire understory of all plant communities at the reserve. Due to the heterogeneity of the landscape, fire intensity was highly variable, with some fire-killed trees standing 10 meters from those that were seemingly unaffected. This heterogeneity provided ideal conditions to observe relationships between *Q. douglasii* mortality, soil microbial population and community structure, and carbon storage.

### 2.2 Tree Selection and Sample Collection

Three dead and three living *Q. douglasii* trees were selected for sampling, based on their accessibility and lack of surrounding woody vegetation. A different tree was sampled on each of 6 different days due to logistical constraints imposed by the sampling method. Tree names, locations, sample dates, and health statuses are summarized in Supplementary Table 1. For each tree, a location within 1m of the tree’s trunk was chosen for sampling and cleared of grass and undecomposed plant litter. A cylindrical soil core (Fat Daddio’s Pro Series SSRD-4030, 100×75 mm) was malleted into the ground, and six successive depth fractions were extracted in a single vertical column (0 - 7.5 cm, 7.5 - 15 cm, 15 - 22.5 cm, 22.5 - 30 cm, 30 cm - 37.5 cm, and 37.5 cm - 45 cm). An access pit was dug alongside the sampling column, and each core was sealed and manually excavated from the side to prevent contamination. Photos illustrating this process can be found in Supplementary Figure 1. Between the removal and handling of each core, all tools were thoroughly washed and sterilized with 70% ethanol. Within 30 minutes of sampling, soil from each core was weighed, homogenized with a 4mm sieve, and subsampled. Each of a 2.0 g and 50 g subsample were flash-frozen on dry ice for nucleic acid and phospholipid-derived fatty acid (PLFA) analysis, respectively. The remaining homogenized soil was stored on water ice for chemical and physical analyses. Within 10 hours, all subsamples for nucleic acid and PLFA analyses were transported to a -70°C freezer, where they were stored until further processing. The remainder of soil samples were refrigerated at 4°C.

### 2.3 Soil Properties

Soil was weighed onto aluminum dishes, allowed to air-dry for 72 hours, weighed again, and finally oven-dried at 105°C overnight and weighed a final time to calculate soil moisture and field-moist, air-dry, and oven-dry conversion factors. Bulk density was calculated by multiplying core volume by moisture-corrected core masses. Soil pH was measured by suspending 5 g field-moist soil in 10.0 mL 0.01 M CaCl_2_, incubating 30 minutes at room temperature, briefly centrifuging, and measuring with an OrionStar A211 pH meter (Thermo Scientific, Waltham, MA). Dry and moist soil colors were measured using a Munsell color book (*Munsell soil color charts*, 2000). Other soil chemical and physical analyses were performed on air-dry soils at Brookside Laboratories, Inc. (New Bremen, OH). These included soil texture (Bouyoucos, 1962), organic matter (Schulte and Hopkins, 1996), Total %C, %N, and C/N ratio (Nelson and Sommers, 1996), Bray 1 Phosphorus (Bray and Kurtz, 1945), cation exchange capacity (Ross and Ketterings, 1995), and Mehlich III Extractable P, Mn, Zn, B, Cu, Fe, Al, S, Ca Mg, K, and Na (Mehlich, 1984). Total carbon storage per depth fraction (reported in metric tons C/hectare) was calculated by multiplying bulk density by %C.

### 2.4 Microbial Population

Microbial biomass (total, bacterial, and fungal) was measured using phospholipid fatty acid analysis, as in Frostegård and Bååth, 1996, by Ward Laboratories, Inc. (Kearney, NE). One 50g soil subsample frozen on dry ice within 30 minutes of sampling was shipped to Ward Labs, where lipids were extracted from frozen soil samples, subjected to alkaline methanolysis, and the resulting fatty acid methyl esters separated and identified using gas chromatography/mass spectrometry. All PLFA (including undifferentiated) were added to calculate total microbial biomass, all bacterial biomarkers were added to calculate all bacterial biomass, and all fungal biomarkers were added to calculate total fungal biomass. While the use and relevance of PLFA for soil community ecology has been challenged (Frostegård et al., 2011), it remains a valid method for measuring overall biomass of broad taxonomic groups (gram +/-bacteria, actinobacteria, and fungi). Potentially over-detailed interpretations (e.g., saprotrophic vs. arbuscular mycorrhizal fungi) were avoided in this study.

### 2.5 Microbial Community

RNA and DNA were dual-extracted from flash-frozen soil samples with the Qiagen RNeasy PowerSoil Total RNA Kit (Qiagen, Carlsbad, CA) and subsequently isolated from each other using the Qiagen RNeasy PowerSoil DNA Elution Kit (Qiagen, Carlsbad, CA). Extraction yield of both RNA and DNA were quantified using a Qubit Fluorometer 4 (Thermo Fisher, Waltham, MA). Any remaining DNA was removed from RNA samples using a RNA Clean & Concentrator Kit (Zymo Research, Irvine, CA). Measurable quantities of both RNA and DNA were successfully extracted from 31 of 36 soil cores. To prepare rRNA for amplification, all extracted RNA samples were reverse transcribed into cDNA following the LunaScript® RT SuperMix Protocol for Reverse Transcription (New England Biolabs, Ipswich, MA). ITS and 16S amplicons were generated in parallel from each DNA and RNA-derived cDNA sample and prepared for metabarcoding according to the two-step PCR workflow presented in Chen et al., 2021, with some modifications. We describe our detailed amplification protocol in Supplementary Material 1. Prepared amplicon libraries were submitted for AVITI (Element Biosciences) sequencing at the UC Davis DNA Technologies Core Laboratory. Due to difficulty amplifying microbial RNA and DNA, especially from deeper soil fractions, we submitted libraries from 21 ITS RNA extractions, 27 16S RNA extractions, and 30 from each of ITS and 16S DNA extractions.

### 2.6 Analysis

#### 2.6.1 Data Management and Basic Statistics

Data was processed using functions written in R (R Core Team, 2024), version 4.4.0, 2024-04-24). Data was cleaned and pre-processed for analyses using functions from the tidyverse package (Wickham et al., 2019). Linear Mixed-Effects Models (LMEs) were computed using the lme4 package in R with tree identity as a random effect (Bates et al., 2015). All LMEs featuring multiple fixed effects were tested with fixed effects as interactive and reported as such if the interaction term was significant; if it was not significant, a model in which fixed effects are additive was reported instead. Response variables (total living fungal biomass, fungi:bacteria ratio, relative abundance of ectomycorrhizal fungi, biomass of ectomycorrhizal fungi) for which initial LMEs did not meet the assumptions of homoscedasticity or normality of residuals were log-transformed before use in final LMEs.

#### 2.6.2 Bioinformatics

Amplicon reads were quality-screened and trimmed of primers using cutadapt (version 4.2 using Python v3.10.8; Martin, 2011), and bioinformatic analyses were performed using DADA2 (Callahan et al., 2016) using default settings, except that reads were quality-filtered using filterandTrim() with minLen = 100. Taxonomy was assigned to quality-filtered, denoised, merged, and chimera-filtered amplicon sequence variants (ASVs) using assignTaxonomy() (ITS: UNITE reference “sh_general_release_dynamic_25.07.2023.fasta” (Nilsson et al., 2019); 16S: SILVA reference “silva_nr99_v138.1_train_set.fa.gz” plus species assignment with addSpecies(), reference “silva_nr99_v138.1_wSpecies_train_set.fa.gz” (Quast et al., 2012). Finally, communities along with their environmental metadata were compiled into phyloseq objects (McMurdie and Holmes, 2013) for further analysis.

#### 2.6.3 Imputed Microbial

*Populations and Traits* Biomass of individual microbial groups (e.g., ectomycorrhizal fungi) was calculated by multiplying the relative abundance of that group represented by the RNA-based amplicon libraries by the overall biomass (ng lipid / g soil) of PLFA for that group (e.g., fungi). Bacterial traits were assigned to 16S ASVs based on their assigned taxonomy by referencing the FAPROTAX database (Louca et al., 2016) using the function X$cal_spe_func() in package microeco (Liu et al., 2021). Fungal traits and niches were assigned to ITS ASVs similarly by referencing the FungalTraits database (Põlme et al., 2020). Percent ectomycorrhizal fungi was specifically determined using the “primary lifestyle” component of FungalTraits.

#### 2.6.4 Microbial Community Statistics

Non-Metric Multidimensional Scaling (NMDS) ordinations and permutational analyses of variance (PERMANOVAs) were calculated using the metaMDS() and adonis2() functions in package vegan (Oksanen et al., 2024). Bray-Curtis dissimilarities were used for all community dissimilarity metrics. Main and interaction effects of depth, tree health, and nucleic acid on microbial community dissimilarity were tested, with tree identity as a blocking factor. Correlations between edaphic data and changes in microbial communities were calculated using function envfit() in vegan; however, many variables were multicollinear, varying with depth. To clarify the relationship between edaphic factors and microbial communities, exploratory factor analysis was performed using function fa() in package psych (Revelle, 2024). Function envfit() was used to fit calculated factors to microbial community and function ordinations.

#### 2.6.5 Ectomycorrhizal-Associated Bacteria

16S ASVs (RNA amplicons only) significantly associated with the biomass of ectomycorrhizal fungi were calculated using functions phyloseq_to_deseq2() and DESeq() in package DeSeq2 (Love et al., 2014). Only RNA-based communities were selected to better represent the living microbial community at the time of sampling (Carini et al., 2016). ASVs were pre-filtered to only include those seen under at least 3 trees, to remove outliers and attempt to control for site-specific effects.

## Results

### 3.1 Sequencing

Other than negative controls, 16S amplicon libraries had an average read depth of 317,578 (36,449 - 1,548,126), while ITS libraries averaged 446,176 reads (80,040 - 2,466,968). An average of 91.4% of 16S reads passed quality control filters, and 96.4% of ITS reads passed. After filtering, read depth for each of two 16S negative controls was 868 and 947, while those for three ITS negative controls were 2,718, 3,010, and 3,284. Rarefaction curves plateaued for all samples between 30k-100k reads (Supplementary Figure 2). We were confident that our communities were represented by more than adequate sequencing depth; to control for the influence of spurious reads seen in our negative controls, we removed reads equal to 0.01% of the lowest read depth for each of 16S (3) and ITS (8) from every ASV in every sample. Where the abundance of any ASV was reduced to 0, it was removed from that sample. Our final read depths for 16S samples ranged from 25,427 to 1,489,955, from 37,687 ASVs; our depths for ITS samples ranged from 73,545 to 2,449,376, from 7,671 ASVs.

### 3.2 Microbial Community Size

Contrary to our predictions, total living microbial biomass was similar in soils associated with living and dead trees (LME, p = 0.503). This biomass was distributed similarly throughout the soil column in both cases (no depth:tree mortality interaction), with less biomass found in deeper soils (p << 0.001, Supplementary Figure 3). Total living fungal and bacterial biomass were also similar between living and dead trees (LMEs, p = 0.07 and p = 0.15, respectively), but in these cases, the distribution of biomass throughout the soil column differed with tree mortality (p = 0.003; p = 0.030; Supplementary Figure 3). Regardless of tree mortality, fungal biomass was much greater in surface soils (0 - 7.5 cm) than at any other depth, but after an initial decrease it plateaued in soils under dead trees while continuing to decline with depth in soils under live trees (Figure 3). This decrease in fungal population size over depth was significant in soils under both living and dead trees (p << 0.001; p = 0.005). Bacterial biomass, however, only decreased significantly over depth in living trees (LME, p < 0.001) and was not correlated with depth in dead trees (LME, p = 0.163). Fungi:bacteria ratio was uncorrelated with tree mortality and negatively correlated with depth (LME, p << 0.001, p = 0.696), with no interaction between the two. On average, bacterial biomass was 1.62x greater than fungal biomass in surface soils (0 - 7.5 cm) but 4.22x greater in our deepest measured soils (37.5 - 40 cm).

### 3.3 Microbial Community Composition

Bacterial and fungal communities were evaluated both in terms of the relative abundance of individual ASVs (“taxonomic composition”) and the relative abundance of discrete functional traits among ASVs (“trait-based” or “functional composition”). Trait data were available (Methods, 2.6.3) for an average of 74.0% (25.5% - 99.9%) of fungal ASVs per sample and 35.7% (20.1% - 65.7%) of bacterial ASVs. Soil depth, tree mortality, and nucleic acid (RNA or DNA) were evaluated as explanatory variables. Soil chemical properties and other edaphic factors were omitted from statistical analyses, as they were highly correlated with each other; a factor analysis (Supplementary Figure 4; Supplementary Table 2) identified two main vectors of covariation among soil chemical and physical properties (Figure 1), which we interpret to represent depth and soil weathering, respectively. PERMANOVAs (Table 1, Figure 1) showed tree mortality to be significantly correlated with both the taxonomic and trait-based composition of fungal communities, but only the trait-based composition of bacterial communities. In all cases, depth correlated significantly with community composition, and changes in composition over depth were distinct between living and dead trees (Table 1). While All correlations above exist independent of the nucleic acid used for community profiling, DNA-based and RNA-based communities generated from the same soil samples were consistently distinct from one another (Table 1, Figure 2). Because communities constructed from amplified rRNA exclusively represent living microbes (Blazewicz et al., 2013; Runte et al., 2021), all subsequent analyses of microbial relative abundance and comparisons to PLFA were performed using RNA-based communities.

**Figure 1.**
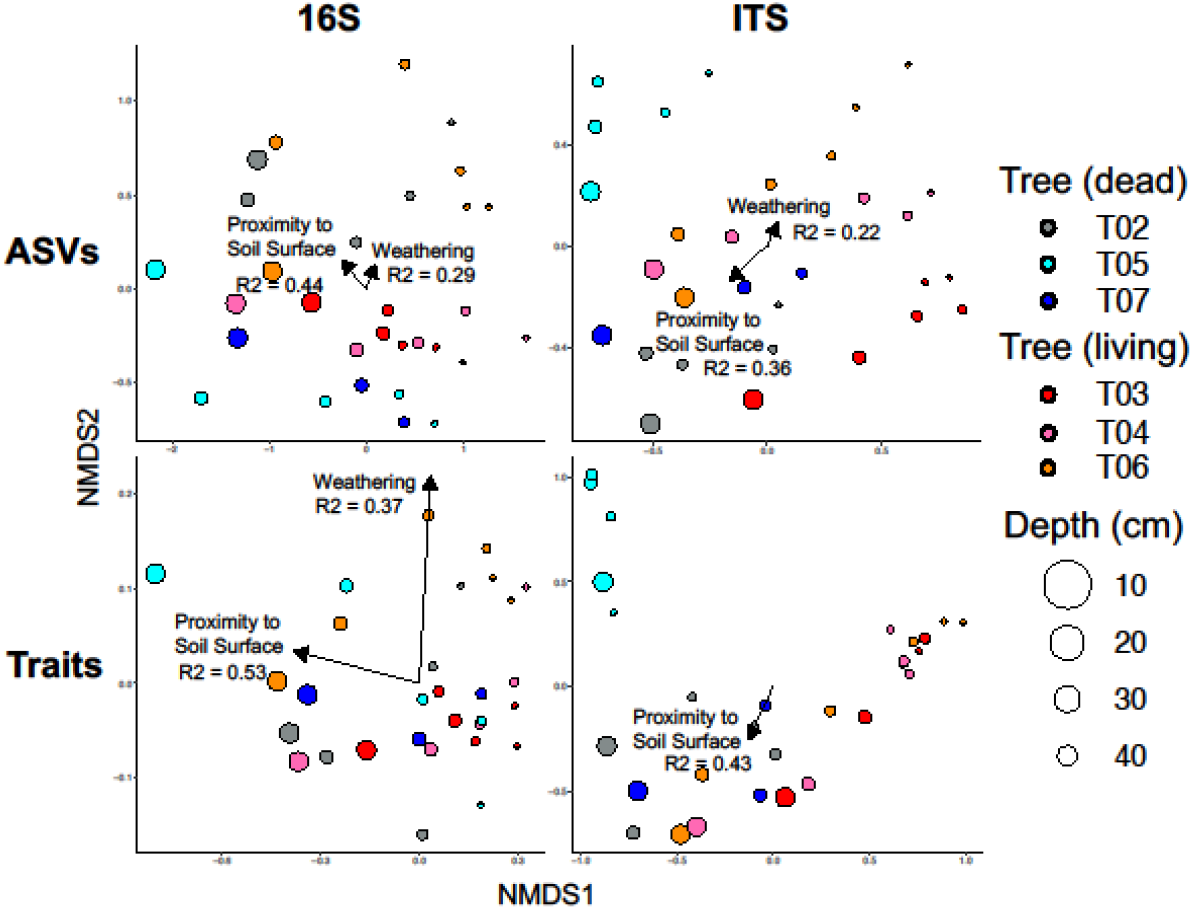
Community dissimilarity in DNA-based (clockwise from top left) 16S taxonomic, ITS taxonomic, ITS trait-based, and 16S trait-based communities. Significant (envfit p < 0.05) factors derived from exploratory factor analysis of edaphic variables are fitted to NMDS ordinations, with R^2^ displayed. Vector length is scaled to R^2^.

**Table 1.** Permutational Analysis of Variance (PERMANOVA) results for 16S and ITS taxonomic (ASVs) and functional (traits assigned using FAPROTAX (16S) and FungalTraits (ITS)) communities. Significance of main or interaction terms (p < 0.05) are shown in bold text. For ITS Traits, there is a significant interaction with nucleic acid; for this test, individual influences of depth and tree health in both RNA- and DNA-based communities are also shown. Degrees of freedom were: 56 for 16S ASVs and Traits; 50 for ITS ASVs and Traits; 20 for ITS RNA-based Traits; 29 for ITS DNA-based Traits.

|  | Factor | R2 | F | Pr(>F) |
| --- | --- | --- | --- | --- |
| **16S ASVs** | **Depth** | **0.17** | **11.50** | **0.001** |
|  | Tree Health | 0.04 | 2.56 | 0.813 |
|  | **Nucleic Acid** | **0.03** | **2.30** | **0.004** |
|  | **Depth x Tree Health** | **0.03** | **2.24** | **0.010** |
|  | Depth x Nucleic Acid | 0.01 | 0.80 | 0.424 |
|  | Tree Health x Nucleic Acid | 0.01 | 0.48 | 0.972 |
| **ITS ASVs** | **Depth** | **0.07** | **4.05** | **0.001** |
|  | **Tree Health** | **0.08** | **4.91** | **0.008** |
|  | **Nucleic Acid** | **0.02** | **1.44** | **0.001** |
|  | **Depth x Tree Health** | **0.04** | **2.05** | **0.001** |
|  | Depth x Nucleic Acid | 0.01 | 0.77 | 0.630 |
|  | Tree Health x Nucleic Acid | 0.02 | 0.93 | 0.135 |
| **16S Traits** | **Depth** | **0.53** | **70.27** | **0.001** |
|  | Tree Health | 0.02 | 2.31 | 0.955 |
|  | **Nucleic Acid** | **0.05** | **5.96** | **0.008** |
|  | **Depth x Tree Health** | **0.03** | **3.36** | **0.050** |
|  | Depth x Nucleic Acid | 0.00 | -0.03 | 0.997 |
|  | Tree Health x Nucleic Acid | 0.01 | 0.72 | 0.456 |
| **ITS Traits** | **Depth** | **0.22** | **22.16** | **0.001** |
|  | **Tree Health** | **0.24** | **25.18** | **0.001** |
|  | **Nucleic Acid** | **0.04** | **4.20** | **0.004** |
|  | **Depth x Tree Health** | **0.04** | **3.90** | **0.004** |
|  | Depth x Nucleic Acid | 0.01 | 0.54 | 0.662 |
|  | **Tree Health x Nucleic Acid** | **0.03** | **3.17** | **0.010** |
| **ITS Traits - RNA** | **Depth** | **0.18** | **7.28** | **0.001** |
|  | **Tree Health** | **0.33** | **13.04** | **0.001** |
|  | Depth x Tree Health | 0.05 | 2.08 | 0.058 |
| **ITS Traits - DNA** | **Depth** | **0.24** | **14.16** | **0.001** |
|  | **Tree Health** | **0.26** | **15.01** | **0.046** |
|  | **Depth x Tree Health** | **0.05** | **2.82** | **0.016** |

**Figure 2.**
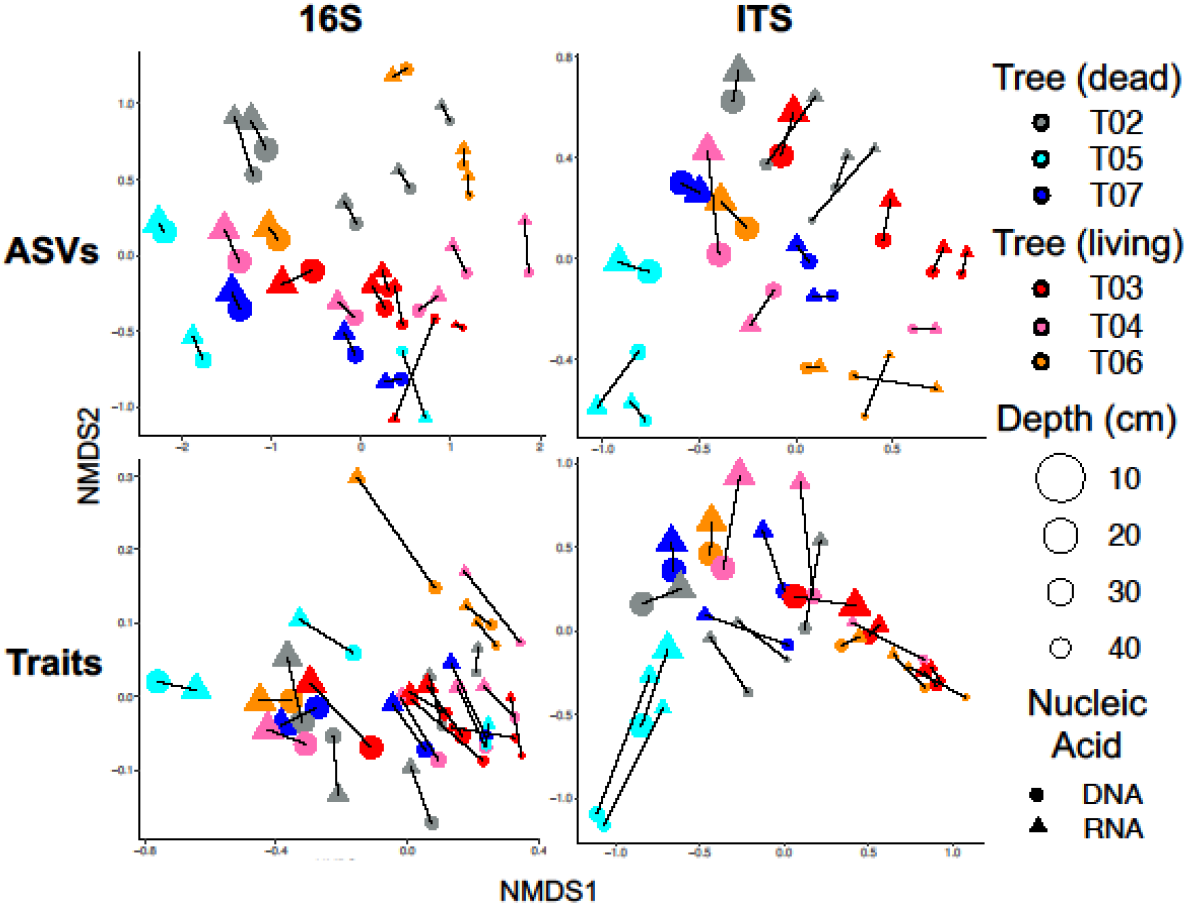
Contrast between RNA-based and DNA-based assessments of (clockwise from top left) 16S taxonomic, ITS taxonomic, ITS trait-based, and 16S trait-based communities. Black bars link RNA-based and DNA-based communities derived from the same soil sample.

### 3.4 Trait-based Composition of Fungal Communities

Due to the low percentage of our bacterial community for which trait data was available through FAPROTAX, further analyses of trait-based bacterial community composition were deemed uninterpretable and have been omitted. Deeper investigation into the trait-based composition of fungal communities, however, revealed stark differences in the prevalence of ectomycorrhizal fungi between soils collected under living and dead trees. The imputed biomass (Methods 2.6.3) of ectomycorrhizal fungi under living trees is consistently greater than that of dead trees (log-transformed LME, p = 0.037, Figure 3); when averaged across depth, soils associated with living trees contained 158 ng of lipids attributed to living ectomycorrhizal biomass per gram of soil, while those associated with dead trees contained only 9 ng / g soil. No significant interaction was found between depth and tree mortality. The log-transformed LME found ectomycorrhizal biomass (lipids) to be positively correlated with depth under both living and dead trees – however, the magnitude of this increase is unlikely to be ecologically relevant (Supplementary Figure 5). Observed changes in ectomycorrhizal biomass over depth were minimal in comparison to the decline in total fungal biomass over depth, which, taken together, correspond to increases in the relative abundance of ectomycorrhizal over depth in soils under both living and dead trees (LME, p << 0.001; log-transformed LME, p = 0.007; Figure 3). Given the overlapping ranges of ectomycorrhizal relative abundance in shallower soils under living and dead trees (Supplementary Figure 5), a combined LME detected no significant correlation between either tree mortality or depth alone to ectomycorrhizal relative abundance – however, a strong interaction effect between the two demonstrates that ectomycorrhizal relative abundance increased more drastically in soils under living trees than dead ones (log-transformed LME, p << 0.001), and separate LMEs confirmed a significant effect of depth in both scenarios (p << 0.001; p = 0.045). Under living trees, ectomycorrhizal fungi accounted for an average of 17.7% of the fungal community in surface soils (0 - 7.5 cm) and an average of 63.9% in deeper soils (below 22.5 cm). This relationship was maintained but diminished under dead trees, where ectomycorrhizal fungi only accounted for an average of 0.1% of fungi in surface soils and 9.1% in deeper soils. While ectomycorrhizal fungi composed a clear majority of fungi in deeper soils under live trees, such dominance was split under dead trees between consumers of plant matter (pathogens and saprotrophs; 49.2%) and fungi for which no trait data was available (43.4%) (Figure 3). Notably, the average ratio of ectomycorrhizal biomass to plant pathogen + saprotroph biomass under live trees was 10.56x, while under dead trees it was 0.23.

**Figure 3.**
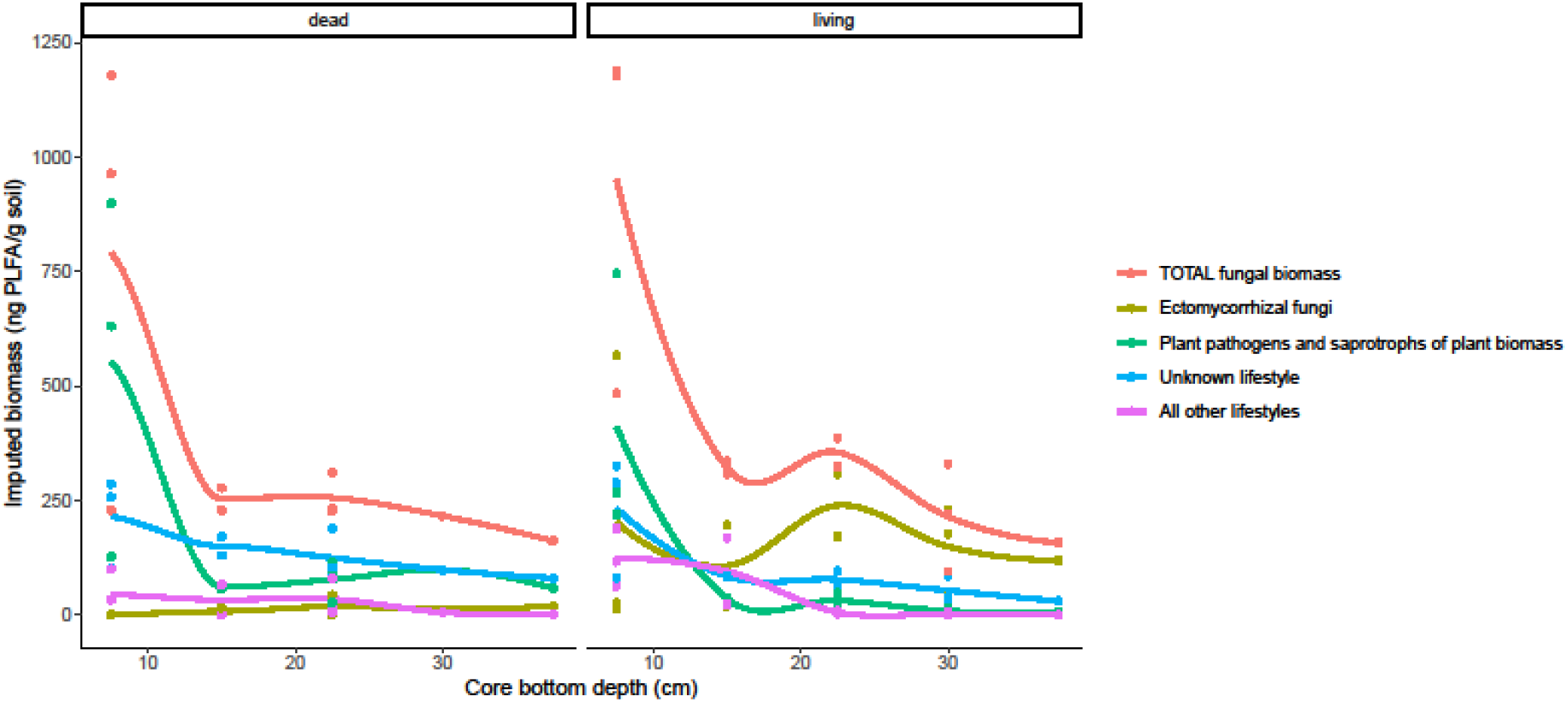
Rolling averages of fungal biomass (ng PLFA / g soil) over depth by functional group. Total fungal biomass is represented in red. All other colors represent the imputed biomass (proportion of fungal community as measured by 16S rRNA metabarcoding * total fungal biomass as measured by PLFA) of discrete functional groups of fungi. Functional groups were assigned to ASVs based on “primary lifestyle” denoted in the FungalTraits database and further agglomerated for visual simplicity. Colored curves are local regressions fitted by applying the “loess” method to geom_smooth() in ggplot2.

### 3.5 Effect of ectomycorrhizae on bacterial community

After removing one high-leverage sample, biomass of ectomycorrhizal fungi was not significantly correlated with bacterial alpha diversity, either observed ASVs or Shannon index. Indicator species analysis showed the relative abundance of 114 bacterial ASVs to be correlated with ectomycorrhizal biomass. Of these, 100 were positively correlated (Supplementary Table 3, Figure 4). The latter “ectomycorrhizal-promoted” bacteria represented up to 23.4% but an average of 6.0% of the active bacterial community in samples collected under live trees. In samples collected under dead trees, these bacteria represented an average of 0.6% of the community (Supplementary Figure 6).

**Figure 4.**
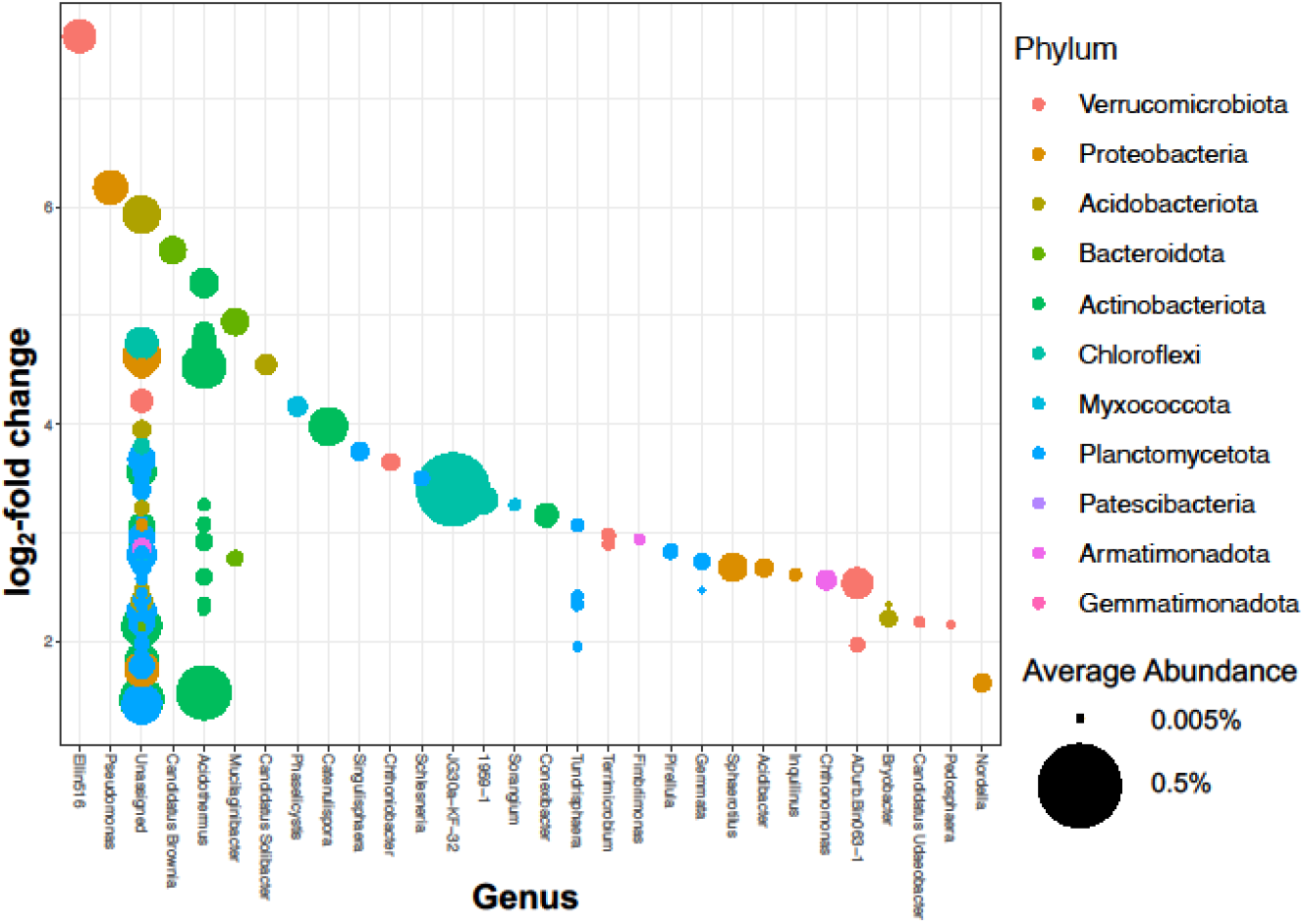
Bacterial taxa whose relative abundance is positively correlated with ectomycorrhizal biomass. Each point represents a bacterial ASV detected in soils associated with at least three trees in our study via RNA-based 16S metabarcoding. Y-axis is log fold change in relative abundance among bacteria per 100 ng lipid / g soil increase in ectomycorrhizal biomass. Abundances are averaged across all samples.

**Figure 5.**
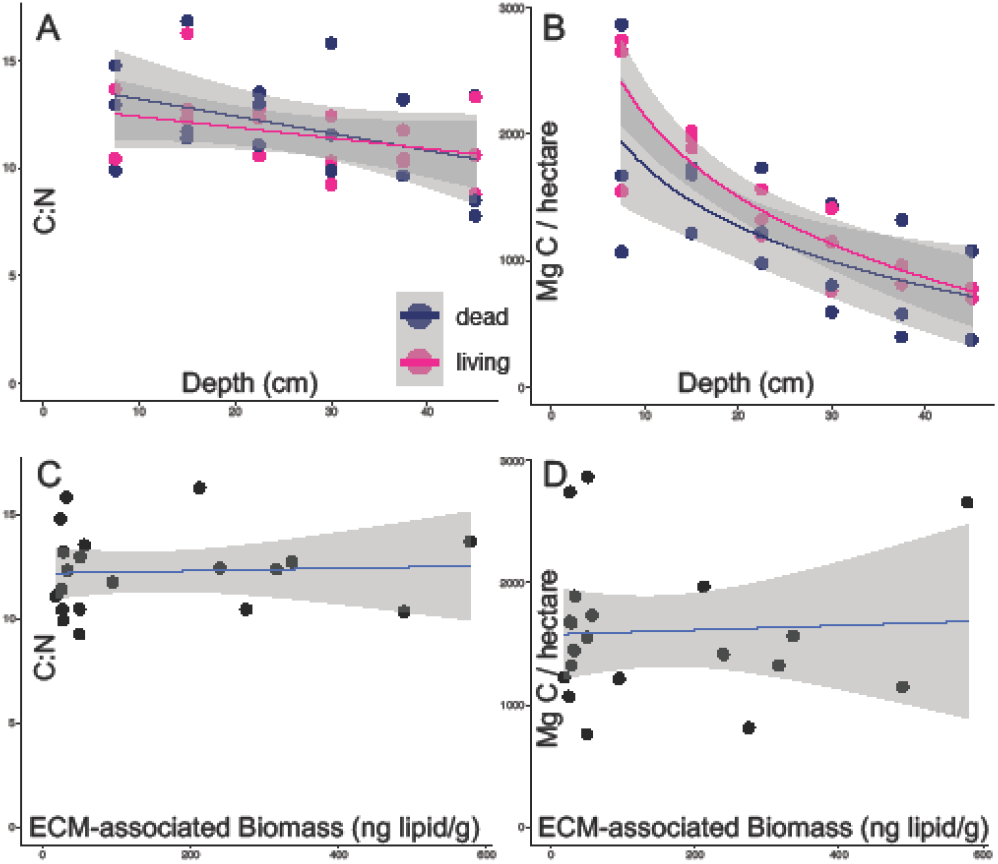
C:N ratio (a, c) and total soil carbon (Mg / ha) (b, d) as related to soil depth, tree mortality, and the biomass sum of ectomycorrhizal fungi and ectomycorrhizal-promoted bacteria (“ECM-associated biomass”). On all panels, best fit lines illustrate linear relationships (A, C, D y ∼ x; B y ∼ log(x)), with gray shading indicating 95% confidence intervals. There are no significant differences between dead and living trees, nor correlations with ectomycorrhizal-associated biomass.

### 3.6 Soil carbon storage

Total soil carbon, adjusted for bulk density and extrapolated to megagrams per hectare, was negatively correlated with depth but not correlated with tree mortality (LME; p << 0.001, p = 0.323). The same was true for C:N ratio (LME; p = 0.002, 0.829). For both response variables, no interaction was found between tree mortality and depth. While total carbon decreased over depth, notable amounts of carbon were recovered from the subsoil; in this dataset, 28.7% of total C is stored below 22.5 cm. To evaluate potential impacts of ectomycorrhizal-mediated carbon flow on soil carbon storage, the biomass sum of ectomycorrhizal fungi and ectomycorrhizal-promoted bacteria was calculated and used as a fixed effect in separate LMEs to total soil carbon and C:N ratio. Contrary to our predictions, no correlation was observed between ectomycorrhizal-promoted biomass and either carbon variable (LMEs; p = 0.960; p = 0.982).

## Discussion

### 4.1 Tree mortality not linked to overall population size of subsoil microbial communities

Contrary to our hypothesis, we did not find that soils associated with living oak individuals contained larger microbial populations than those associated with dead individuals. This result is unsurprising in oak savanna surface soils, where roots of understory plants and nearby leaf litter may also provide carbon substrate to microbes, and reflects an existing study which sampled to a maximum depth of 10 cm (García-Angulo et al., 2020). It is more surprising that this was also true in subsoils, where oak roots are often the only living plants able to deposit new carbon. The microbial populations we observed under dead oaks may still be supported by carbon substrate deposited by the tree before its death; while the trees in our study are assumed to have died only three years prior to soil sampling, legacy effects of rhizodeposited carbon on microbial population size have been observed millenia after the death of the root (Gocke et al., 2017). Within the scope of this study, we cannot separate the effects of historically rhizodeposited carbon substrates and active carbon imports on microbial population size.

### 4.2 Tree mortality transforms the trait-based composition of subsoil fungal communities

Mortality-correlated compositional divergence was particularly drastic in trait-based fungal communities (Figure 1). Despite notable variation among our three living trees in their overall average relative abundance of ectomycorrhizal fungi (T03 = 67%, T04 = 18%, T06 = 50%, Supplementary Figure 7), ectomycorrhizae dominate the fungal niche in subsoils associated with living trees, but disappear almost entirely in subsoils associated with dead trees (Figure 3). Ectomycorrhizal fungi composed the single most abundant fungal guild observed in all living-tree associated soils deeper than 22.5 cm, and represented a clear majority of fungi in T03 and T06 (Supplementary Figure 7). In T04, for which rRNA was only successfully extracted from a single core below 22.5cm (22.5 cm - 30 cm), ectomycorrhizal fungi still represented 43% of the fungal community; 40% of fungi in this core had no assigned function. In association with dead trees, ectomycorrhizal relative abundances were < 20% at comparable depths in all trees and < 10% in two out of three. This implies that within the three years following the death of these three oaks, ectomycorrhizae largely vacated the subsoil fungal niche; in their stead, we observe saprotrophic and plant-pathogenic fungi composing just under half of the fungal community, with much of the remainder functionally unaccounted for (Figure 3). Given that we find no evidence of the fungal niche expanding or contracting (i.e., total fungal biomass increasing or decreasing) with tree mortality over the time scale of this study, we can infer that these shifts correspond to a loss of ectomycorrhizal biomass and an increase in pathogen/saprotroph biomass in the wake of tree mortality, as visualized in Figure 3. These patterns could be explained by the Gadgil effect (Gadgil and Gadgil, 1975; Fernandez and Kennedy, 2016), which posits that ectomycorrhizal fungi compete with fungal saprotrophs for water and mineral resources, reducing populations and activity of saprotrophs. The Gadgil effect further suggests that ectomycorrhizae-saprotroph competition results in reduced decomposition rates of SOM, though a recent review found evidence of this to be inconsistent and factors modulating this outcome are unknown (Fernandez and Kennedy, 2016).

### 4.3 Tree mortality structures spatial distribution of fungi, bacteria over depth

While we expected to see differential patterns in the distribution of fungal and bacterial biomass over depth between living and dead trees, the patterns we observed opposed the ones predicted; for both fungi and bacteria, population sizes decrease more consistently over depth in soils associated with living trees than dead trees. While depth-agnostic evaluations of bacterial and fungal biomass find no difference between living and dead trees, the maximums and minimums are more extreme under living trees than dead ones. At the surface, the heightened fungal biomass maximum may be explained by SOM inputs from the recent deposition of oak leaf litter (García-Angulo et al., 2020) – given that dead trees in this study were assumedly killed by wildfire, surface organic matter which burned off during that event may not have been replaced by new leaf litter underneath them as it likely has been replaced under living trees over the past three years. As a source of both moisture and nitrogen, more abundant SOM could lead to a diminished magnitude of Gadgil effect competition, which could explain the simultaneously sizable living-tree populations of ectomycorrhizal fungi and saprotrophs in the topmost core alone (Figure 3). At the surface, these groups appear to contribute additively to the overall fungal population under living trees, such that the relative absence of active ectomycorrhizal fungi under dead trees could explain lesser overall fungal biomass in surface soils. In subsurface soils, however, the replacement of saprotrophs and pathogens by ectomycorrhizal fungi is near complete; therefore, we speculate that different patterns of overall fungal biomass decline in living trees and dead trees might reflect greater depth sensitivity among the ectomycorrhizal fungi in this ecosystem than members of the agglomerated pathogen/saprotroph group. In bacterial populations, the obscurity of ecological roles among the majority of our detected ASVs limit such speculation.

### 4.4 Living biomass of ectomycorrhizae is correlated with altered bacterial community

Our observation that increased biomass of ectomycorrhizal fungi is correlated with changes in the relative abundance of numerous bacterial ASVs in field soils corroborates existing experimental evidence that mycorrhizae shape their local bacterial communities. Most research on mycorrhizal influence over bacterial community composition has focused on arbuscular mycorrhizal fungi (Kakouridis et al., 2024), with only a single experimental study demonstrating this phenomenon in ectomycorrhizal fungi (Berrios et al., 2024). Unlike (Berrios et al., 2024), we found no significant effects of ectomycorrhizal fungi on bacterial species alpha diversity. The bacterial genera we observed to be correlated with ectomycorrhizal fungi and the vector of our correlations were also largely incongruous with (Berrios et al., 2024); several genera (*Conexibacter, Chtoniobacter, Reyranella*) were less abundant in ectomycorrhizal fungi + treatments there but positively correlated with ectomycorrhizal biomass in our dataset, and only a single genus (*Tundrisphaera*) was positively associated with ectomycorrhizal fungi in both. Disagreement between these two studies might be coarsely attributed to the different species of ectomycorrhizal fungi and plant hosts involved, as Berrios et al., 2024 focused on *Pinus muricata* D.Don (Bishop pine) colonized by *Suillis pungens*, neither of which appear in oak savannas.

While trait data was unavailable through FAPROTAX for most of our ectomycorrhizal-associated bacterial ASVs, a literature review of our five most intensely ectomycorrhizal-promoted bacterial genera reveal that three of them (*Acidothermus, Candidatus Solibacter, Pseudomonas*) have constituent species with the genetic capacity for chitin-degradation (Barabote et al., 2009; Ward et al., 2009; Lalucat et al., 2022). Genes which code for glycoside hydrolases are generally conserved at the genus level in bacteria (Martiny et al., 2015), so we infer that the *Acidothermus* and *Candidatus Solibacter* ASVs observed in this study share this capability. The *Pseudomonas* genus is particularly speciose and contains non-fungivorous specialists as well as fungivores (Lalucat et al., 2022). *Acidothermus*, notably, is the most abundant named genus among bacterial ASVs which were positively associated with ectomycorrhizal fungi in our dataset, and the annotated genome of its type species has a similar number of genes for the degradation of chitin as it does for xylan and cellulose (Barabote et al., 2009). In AMF, the promotion of *Candidatus Solibacter, Pseudomonas*, and several other bacterial clades by mycorrhizae has been linked to the consumption of mycorrhizal carbon by their constituents, and to the subsequent conversion of 25% of the net mycorrhizal C soil input into MAOM over a 6-week period (Kakouridis et al., 2024). Our results suggest that a similar pattern of C flow is possible in our system, and could be better illustrated in future field studies by the use of metatranscriptomics, in controlled companion experiments via 13C isotope labeling, and in both scenarios through SOM fractionation. Without these techniques, however, we cannot determine the role of ectomycorrhizal-associated bacteria in the microbial food chain of our study system – especially as the aforementioned genera all contain versatile saprotrophs, and other heavily promoted genera (*Mucilaginibacter, Candidatus Brownia* - an insect endosymbiont) have no documented chitinolytic capabilities (Pankratov et al., 2007; Gruwell et al., 2010). High variability in the combined relative abundance of ectomycorrhizae-associated bacteria between living oak individuals further obscures their role; while T03 and T06 harbored similar amounts of ectomycorrhizal fungi, ectomycorrhizal-promoted bacteria were over 2x more abundant in T06 than T03 (Supplementary Figure 6). Despite uncertainty surrounding these ectomycorrhizal-associated taxa and the unknown prevalence of fungivorous bacteria, the dominance of ectomycorrhizal fungi among subsoil fungal biomass under live trees may still determine the origin of carbon assimilated by generalist fungivore bacteria living at depth. Conversely, in soils associated with dead trees, at least half of the substrate bacterial fungivores have access to come from saprotrophs and pathogens decomposing plant matter, and is therefore associated with the degradation of existing SOM.

### 4.5 Tree health and ectomycorrhizal-associated biomass do not correlate with soil carbon storage

While we did not observe differences in total carbon storage between living and dead trees, we did observe notable amounts of carbon in soils below 22.5cm. Given that we expected subsoils associated with living oaks to store more carbon as a result of harboring larger microbial populations, the lack of correlation between tree mortality and soil carbon storage is unsurprising in light of the lack correlation between mortality and total microbial biomass. As with total microbial biomass, however, the similarity in total stored carbon between living and dead trees in our study may be a result of the recency of death. Over time, the cessation of new C inputs by dead oaks and the already distinct microbial communities associated with them may result in divergent consequences for carbon cycling and soil health. As described in *4.2*, the potential suppression by ectomycorrhizal fungi of SOM decomposition by fungal saprotrophs could increase SOM stability under living trees and throughout the ectomycorrhizal hyposphere. These effects may be obscured in this dataset by the local abundance of historically rhizodeposited carbon under dead trees, as our soil sampling strategy did not discriminate between rhizosphere, hyposphere, and bulk soil.

## Conclusion

While we found no evidence that tree death in oak savannas results in a local net loss of soil carbon or soil microbial biomass within a three-year period, we present evidence that it does result in altered distributions of soil fungal and bacterial biomass over depth. This is accompanied by shifts in microbial community composition, many of which are most evident below 20 centimeters. Shifts in functional composition of soil fungi are particularly drastic, and are driven by the dominance of ectomycorrhizal fungi in subsoils under living trees. Studies of tree root-associated microbes must sample to depths of at least 30 centimeters to observe the most drastic effects of tree activity on these microbiomes, and by extension, their soil habitat. We also demonstrate important differences between DNA-based and rRNA-based soil microbial community assembly in this ecosystem and advocate for the use of rRNA metabarcoding to evaluate contemporary soil microbiomes. We further present evidence that, in accordance with investigations of tree effects on subsoil in other ecosystems (Beule et al., 2022), the structure of microbial communities of savanna subsoils is heavily influenced by both living oak trees and the ectomycorrhizal fungi they host. We did not, however, find evidence that ectomycorrhizal fungi or their associated bacteria promote carbon storage in this ecosystem. In order to better understand the impact of ectomycorrhizal trees and ectomycorrhizal-dispersed carbon on subsoil ecosystems, future studies should combine depth-sensitive soil sampling with metatranscriptomic analyses of soil microbial communities. There is a large pool of carbon in oak savannas below 20 centimeters, and we have demonstrated extreme shifts in the functional profile of microbial communities interacting with this carbon only three years after sudden tree mortality. To predict and control the fate of this carbon, researchers must deeply and specifically explore the mechanisms of these interactions in arid savanna landscapes, in which sudden mass oak mortality events are expected to increase in frequency and severity.

## Acknowledgements

We are grateful to Demorie Galarza, Brittany Long, Dustin Lower, and Rosalyn Muñiz for their work collecting field samples, and to Ross Brennan for facilitating access to Quail Ridge.

## Author contributions

Anna Goodman: Data Curation, Writing - Original draft, Writing - review & editing, Visualization Ernest Walker: Investigation, Data Curation, Writing - review & editing

Glade Bogar: Conceptualization, Investigation, Data Curation, Writing - Original draft, Writing - review & editing, Visualization, Project administration

Laura Bogar: Conceptualization, Investigation, Writing - review & editing, Supervision, Project administration

