## Supplementary Figure 2 for "Functional composition of subsoil microbial communities changes with oak mortality"

Supplementary Figure 2. Rarefaction curves for 16S and ITS after quality-control trimming (removed 3 reads from all 16S ASVs, 8 reads from all ITS ASVs).

16S:

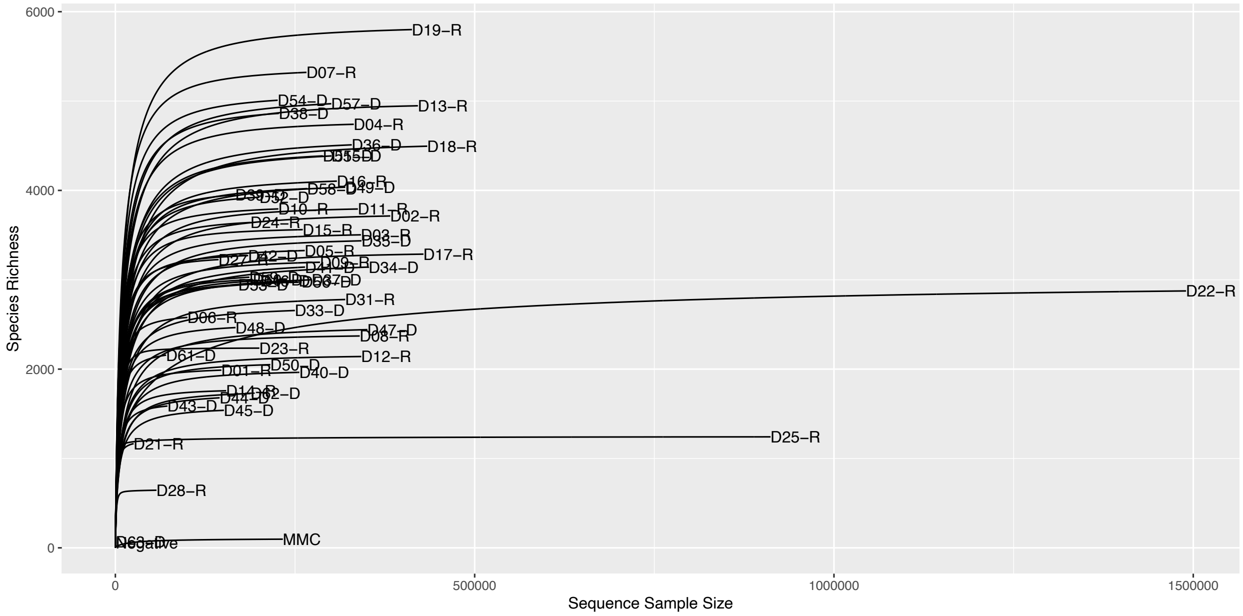


ITS:

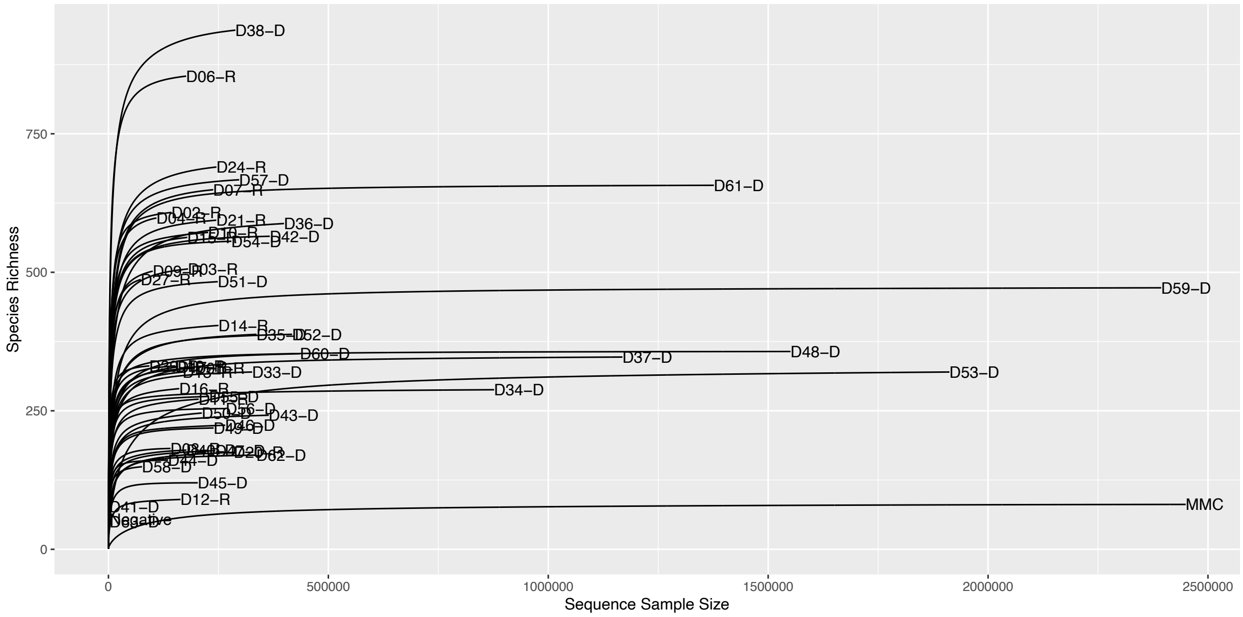
