## Supplementary Figure 3 for "Functional composition of subsoil microbial communities changes with oak mortality"

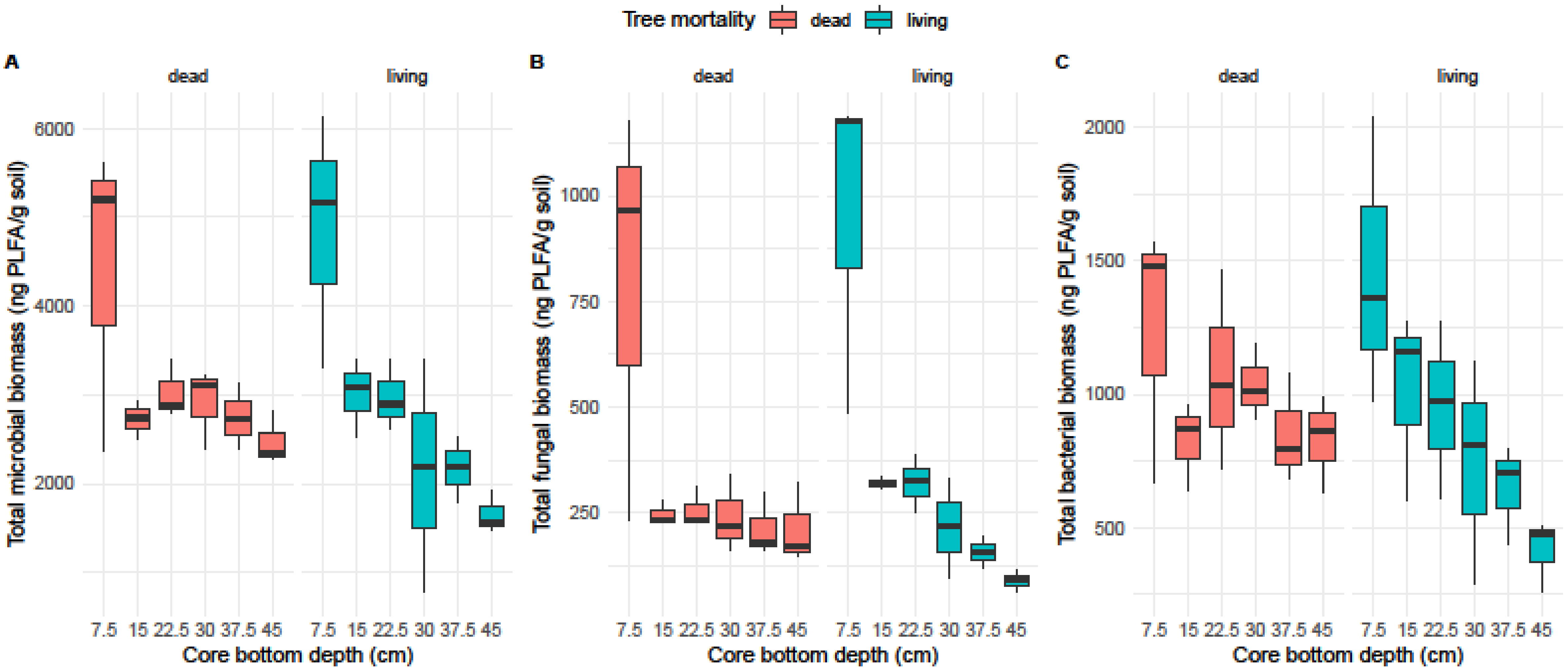


Supplementary Figure 3. Box and whisker plots showing the distribution of fungal (B), bacterial (C), and total (A) microbial biomass across discrete soil depths. Biomass is represented by g PLFA / g soil. Box boundaries are the first and third quartiles, and whiskers extend to extreme values within 1.5 x the interquartile range. Horizontal lines represent the median value. Data from soils collected under dead trees are represented in red, while data from soils collected under living trees are represented in blue; n = 3 within each box.
