## Supplementary Figure 5 for "Functional composition of subsoil microbial communities changes with oak mortality"

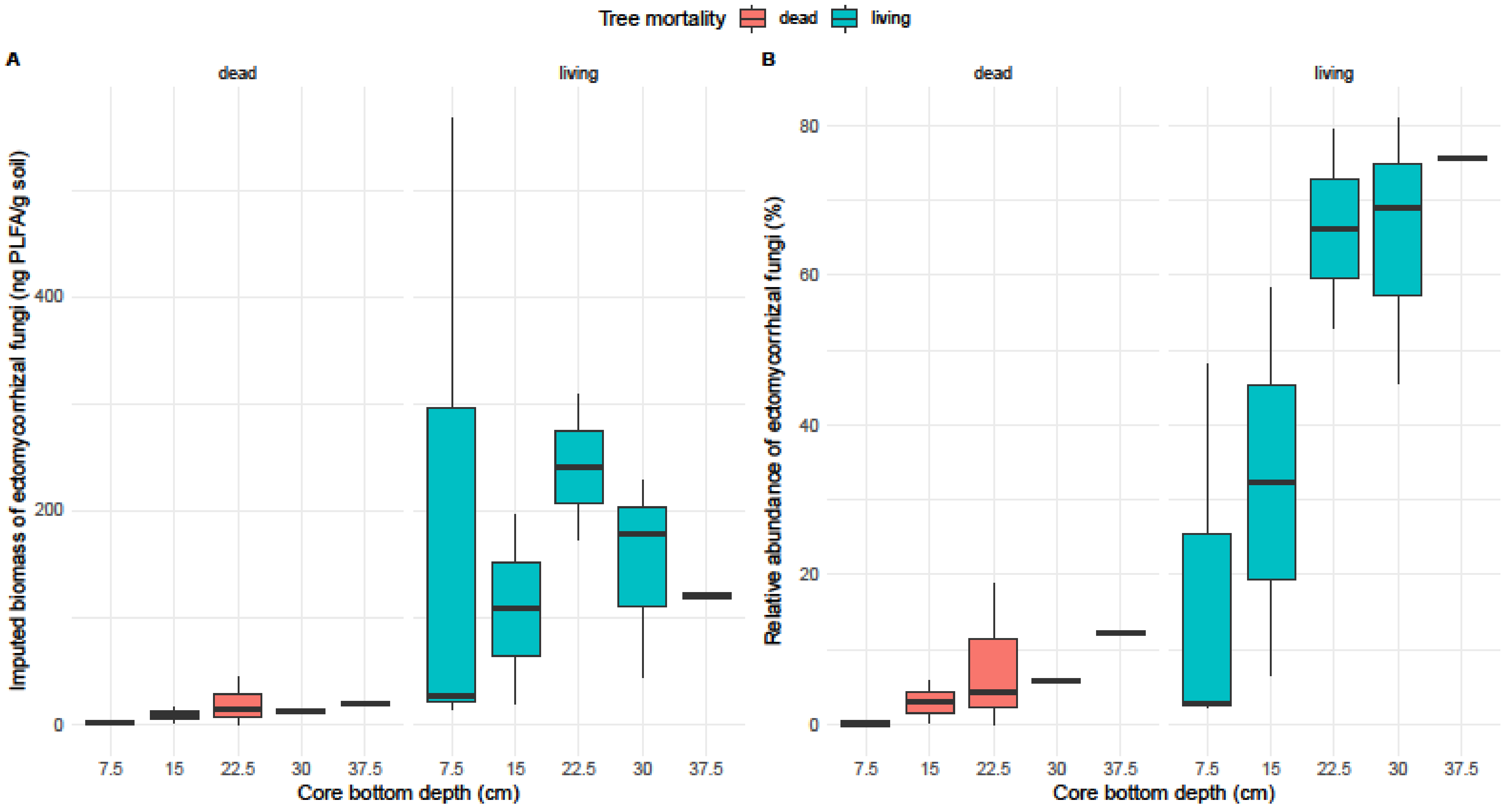


Supplementary Figure 5. Box and whisker plots showing the distribution of imputed biomass (A) and relative abundance (B) of ectomycorrhizal fungi across discrete soil depths. Imputed biomass was calculated as: the proportion of the fungal community (as measured by 16S rRNA metabarcoding) assigned to the ectomycorrhizal primary lifestyle * total living fungal biomass as measured by PLFA. Relative abundance was calculated by multiplying the aforementioned proportion by 100. Box boundaries are the first and third quartiles, and whiskers extend to extreme values within 1.5 * the interquartile range. Horizontal lines represent the median value. Data from soils collected under dead trees are represented in red, while data from soils collected under living trees are represented in blue. Sample size is variable between boxes due to complications with RNA extraction and amplification for metabarcoding. In “dead” panels, from left to right, n = 3, 2, 3, 2, 1. In “living” panels, n = 3, 2, 2, 3, 3.
