## Supplementary Figure 4 for "Functional composition of subsoil microbial communities changes with oak mortality"

Supplementary Figure 4. Correlations between soil edaphic variables and calculated factors made in exploratory factor analysis.

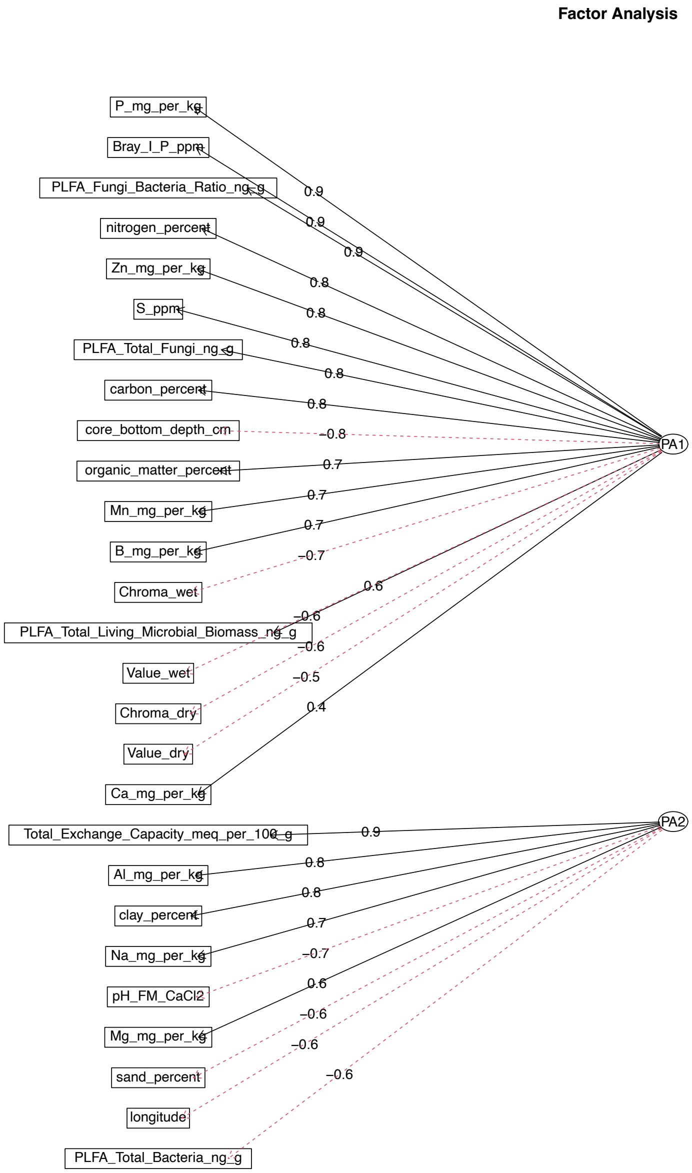
