## Supplementary Figure 6 for "Functional composition of subsoil microbial communities changes with oak mortality"

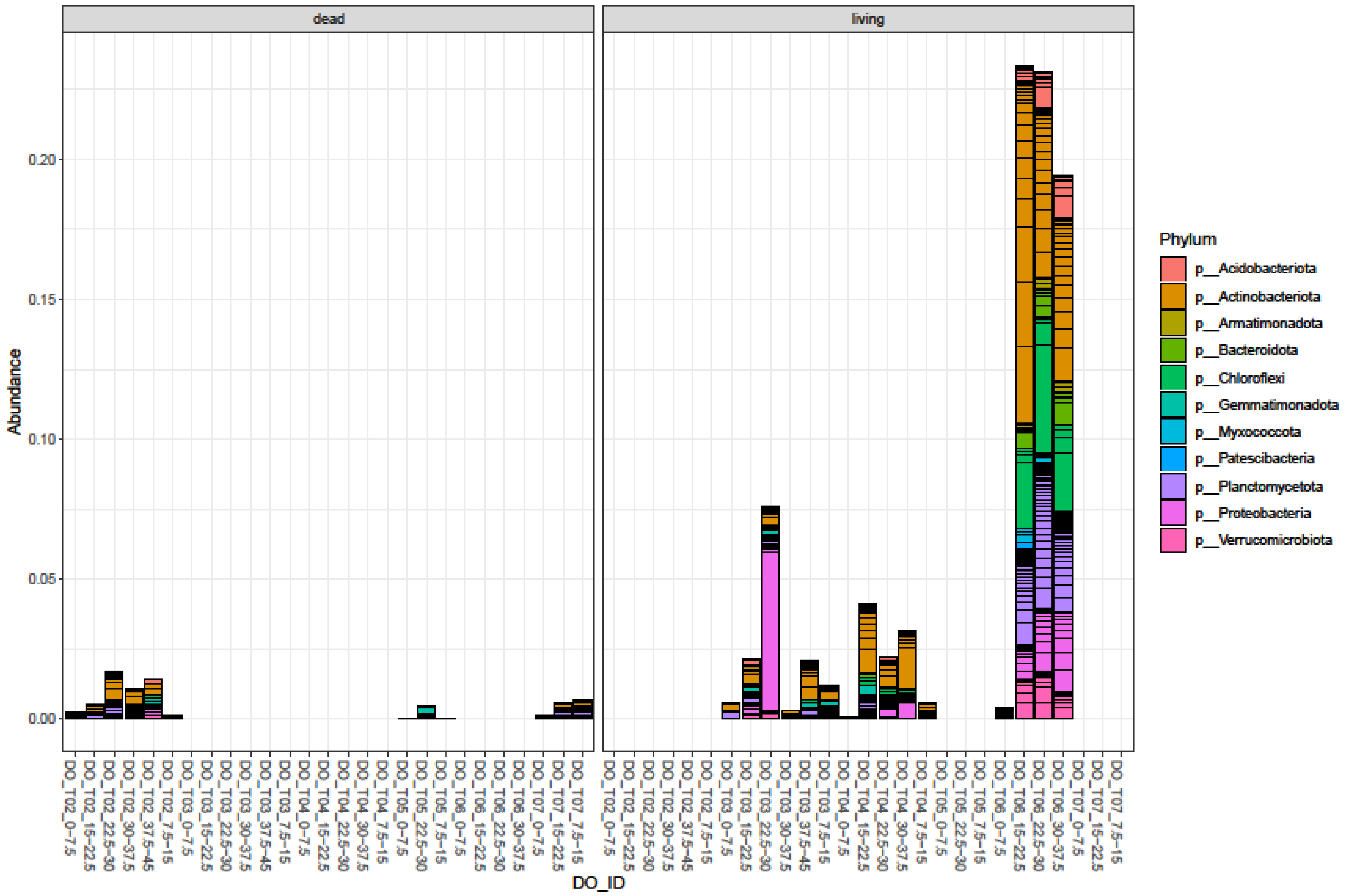


Supplementary Figure 6. Relative abundance of ectomycorrhizal-promoted ASVs within whole-sample bacterial communities as measured by 16S rRNA metabarcoding. Y-axis represents the proportion (0-1) of the RNA-based bacterial community within each discrete soil sample. On the X-axis, soil sample names denote both tree identity and soil depth in cm, and are clustered by tree identity. ASVs are colored according to their genus.
