## Supplementary Material 1 for "Functional composition of subsoil microbial communities changes with oak mortality"

**Supplementary Material 1: Detailed Amplification Protocol:**

We modified a two-step PCR workflow (Chen et al., 2021) to prepare 16S and ITS amplicon libraries for sequencing using the following materials and methods. This protocol features a low-cycle 1st step amplification followed by a low-cycle 2nd step barcode amplification, to limit the overall number of PCR cycles to the barest minimum, to reduce amplification bias and artifacts in amplicon libraries.

Bacterial 16S was amplified with 515f-806r primers (Caporaso et al., 2011) and fungal ITS was amplified with 5.8S-ITS4 primers (Chen et al., 2021), both with equimolar concentrations of primers with six different frame-shift sequences added (final concentration 10 uM) to decrease sequencing error and reduce the need for PhiX. The primers from Chen et al. (2021) were modified to use modern Nextera annealing sites in both the 5’ and 3’ directions.

Our primer sequences were as follows:

| Name | Sequence 5'->3' (Nextera annealing site + frameshift + linker + gene-specific primer) |
| --- | --- |
| 5.8S_Fun_f1 | TCGTCGGCAGCGTCAGATGTGTATAAGAGACAGNNNNNNNN AG AACTTTYRRCAAYGGATCWCT |
| 5.8S_Fun_f2 | TCGTCGGCAGCGTCAGATGTGTATAAGAGACAGNNNNTNNNN AG AACTTTYRRCAAYGGATCWCT |
| 5.8S_Fun_f3 | TCGTCGGCAGCGTCAGATGTGTATAAGAGACAGNNNNCTNNNN AG AACTTTYRRCAAYGGATCWCT |
| 5.8S_Fun_f4 | TCGTCGGCAGCGTCAGATGTGTATAAGAGACAGNNNNACTNNNN AG AACTTTYRRCAAYGGATCWCT |
| 5.8S_Fun_f5 | TCGTCGGCAGCGTCAGATGTGTATAAGAGACAGNNNNGACTNNNN AG AACTTTYRRCAAYGGATCWCT |
| 5.8S_Fun_f6 | TCGTCGGCAGCGTCAGATGTGTATAAGAGACAGNNNNTGACTNNNN AG AACTTTYRRCAAYGGATCWCT |

| ITS4_Fun_f1 | GTCTCGTGGGCTCGGAGATGTGTATAAGAGACAGNNNNN AG AGCCTCCGCTTATTGATATGCTTAART |
| --- | --- |
| ITS4_Fun_f2 | GTCTCGTGGGCTCGGAGATGTGTATAAGAGACAGNNTNNN AG AGCCTCCGCTTATTGATATGCTTAART |
| ITS4_Fun_f3 | GTCTCGTGGGCTCGGAGATGTGTATAAGAGACAGNNCTNNN AG AGCCTCCGCTTATTGATATGCTTAART |
| ITS4_Fun_f4 | GTCTCGTGGGCTCGGAGATGTGTATAAGAGACAGNNACTNNN AG AGCCTCCGCTTATTGATATGCTTAART |
| ITS4_Fun_f5 | GTCTCGTGGGCTCGGAGATGTGTATAAGAGACAGNNGACTNNN AG AGCCTCCGCTTATTGATATGCTTAART |
| ITS4_Fun_f6 | GTCTCGTGGGCTCGGAGATGTGTATAAGAGACAGNNTGACTNNN AG AGCCTCCGCTTATTGATATGCTTAART |

| Name | Sequence 5'->3' |
| --- | --- |
| 515F_f1 | TCGTCGGCAGCGTCAGATGTGTATAAGAGACAGNNNNNNNNGAGTGCCAGCMGCCGCGGTAA |
| 515F_f2 | TCGTCGGCAGCGTCAGATGTGTATAAGAGACAGNNNNTNNNN GA GTGCCAGCMGCCGCGGTAA |
| 515F_f3 | TCGTCGGCAGCGTCAGATGTGTATAAGAGACAGNNNNCTNNNN GA GTGCCAGCMGCCGCGGTAA |
| 515F_f4 | TCGTCGGCAGCGTCAGATGTGTATAAGAGACAGNNNNACTNNNN GA GTGCCAGCMGCCGCGGTAA |
| 515F_f5 | TCGTCGGCAGCGTCAGATGTGTATAAGAGACAGNNNNGACTNNNN GA GTGCCAGCMGCCGCGGTAA |
| 515F_f6 | TCGTCGGCAGCGTCAGATGTGTATAAGAGACAGNNNNTGACTNNNN GA GTGCCAGCMGCCGCGGTAA |

| 806R_f1 | GTCTCGTGGGCTCGGAGATGTGTATAAGAGACAG NNNNN AC GGACTACHVGGGTWTCTAAT |
| --- | --- |
| 806R_f2 | GTCTCGTGGGCTCGGAGATGTGTATAAGAGACAGNNTNNN AC GGACTACHVGGGTWTCTAAT |
| 806R_f3 | GTCTCGTGGGCTCGGAGATGTGTATAAGAGACAGNNCTNNN AC GGACTACHVGGGTWTCTAAT |
| 806R_f4 | GTCTCGTGGGCTCGGAGATGTGTATAAGAGACAGNNACTNNN AC GGACTACHVGGGTWTCTAAT |
| 806R_f5 | GTCTCGTGGGCTCGGAGATGTGTATAAGAGACAGNNGACTNNN AC GGACTACHVGGGTWTCTAAT |
| 806R_f6 | GTCTCGTGGGCTCGGAGATGTGTATAAGAGACAGNNTGACTNNN AC GGACTACHVGGGTWTCTAAT |

1st-step PCR reactions were as follows:

| **Reagent** | **uL/rxn** |
| --- | --- |
| NEB Q5 Hi-Fi PCR Master Mix (2X) | 5.0 |
| Forward Primers (TS4_Fun_f1-f6 or 515_f1-f6) | 0.25 |
| Reverse Primers (5.8S_Fun_f1-f6 or 806_f1-f6) | 0.25 |
| Water | 6.0 |
| Template | 0.5 |
| **TOTAL** | **12.0** |

1st-step thermocycler settings were as follows:

| **Cycles** | **Temp (degrees C)** | **Time (minutes:seconds)** | **Ramp** |
| --- | --- | --- | --- |
| x1 | 98 | 10:00 |  |
| x15 | 98 | 1:00 |  |
|  | 55 | 2:00 | -0.3C/cycle |
|  | 72 | 2:00 |  |
| x1 | 72 | 10:00 |  |

1st-step amplifications were cleaned with Cytiva Sera-mag Select (Cytiva, Marlborough, MA) magnetic beads at a 0.8x sample ratio.

Note that after 15 cycles, amplicons are at too low a concentration to detect by gel electrophoresis or fluorometer. Therefore, no quality check is done between 1st and 2nd step amplifications.

2nd-step PCR reactions added unique dual-indexed barcode pairs, rather than single-index barcodes as in Chen et al., 2021. PCR reactions were as follows:

| **Reagent** | **uL/rxn** |
| --- | --- |
| NEB Q5 Hi-Fi PCR Master Mix (2X) | 10.0 |
| Forward Unique Barcode (5/6 uM) | 6.0 |
| Reverse Unique Barcode (5/6 uM) | 6.0 |
| Water | 1.0 |
| Template | 2.0 |
| **TOTAL** | **25.0** |

2nd-step thermocycler settings were as follows:

| Cycles | Temp (degrees C) | Time (minutes:seconds) | Ramp |
| --- | --- | --- | --- |
| x1 | 94 | 10:00 |  |
| x15 | 94 | 1:00 |  |
|  | 62.5 | 2:00 | -0.3C/cycle |
|  | 72 | 2:00 |  |
| x1 | 72 | 10:00 |  |

2nd-step amplification products were checked by gel electrophoresis, and amplification was repeated where amplifications failed or unexpected bands were seen. Successful amplifications were cleaned again with Cytiva Sera-mag Select magnetic beads, now at a 0.7x sample ratio. These cleaned products were checked by gel electrophoresis a final time, then quantified by Qubit Fluorometer 4 (Thermo Fisher, Waltham, MA), pooled in equimolar concentrations, and sent for sequencing.
