## Supplementary Table 1 for "Functional composition of subsoil microbial communities changes with oak mortality"

| Tree | Latitude | Longitude | Sample Date | Health |
|---|---|---|---|---|
| T02 | 38.48673 | -122.15038 | 08 May 2023 | dead |
| T03 | 38.486593 | -122.150304 | 20 May 2023 | living |
| T04 | 38.488092 | -122.150914 | 22 May 2023 | living |
| T05 | 38.488087 | -122.151009 | 26 May 2023 | dead |
| T06 | 38.488087 | -122.151882 | 28 May 2023 | living |
| T07 | 38.488857 | -122.151742 | 29 May 2023 | dead |
