## Supplementary Figure 1 for "Functional composition of subsoil microbial communities changes with oak mortality"

Supplementary Figure 1. Photos illustrating the depth-resolved soil sampling method described in Methods 2.2. Photos are arranged to represent the method chronologically from left to right, but each shows the sampling area under a different tree (labeled below). In the second row, a photo of Tree 04 is annotated to show the location of the sampling area for spatial context.

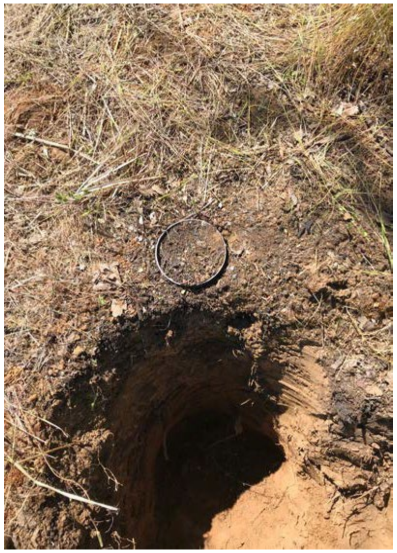
Tree 03

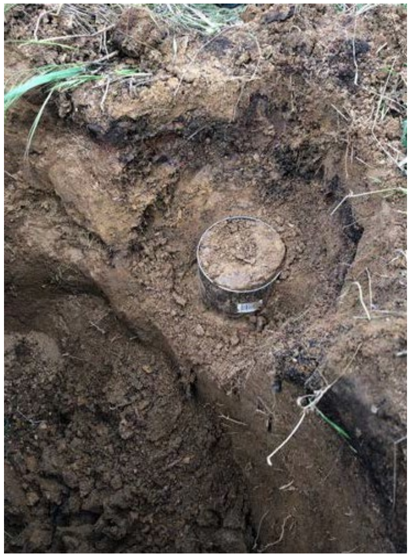
Tree 02

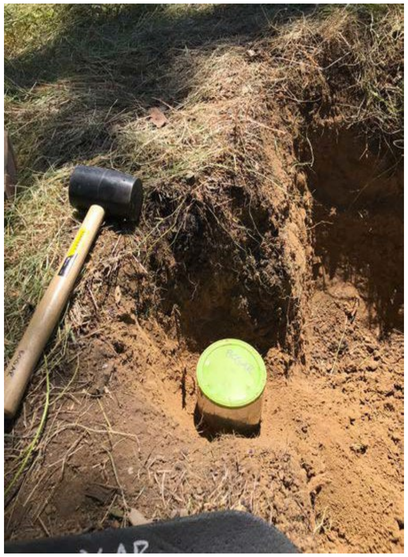
Tree 04

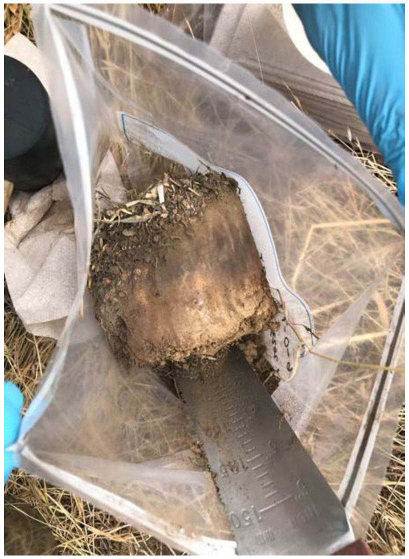
Tree 06

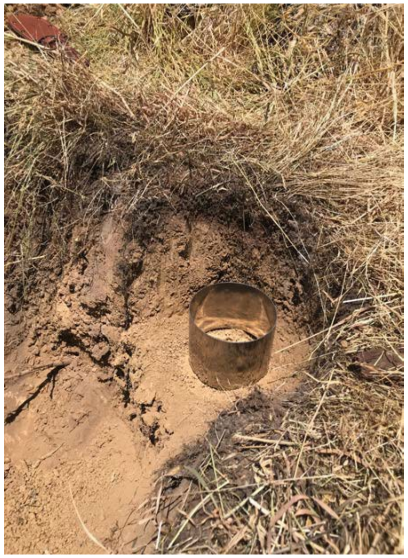
Tree 05

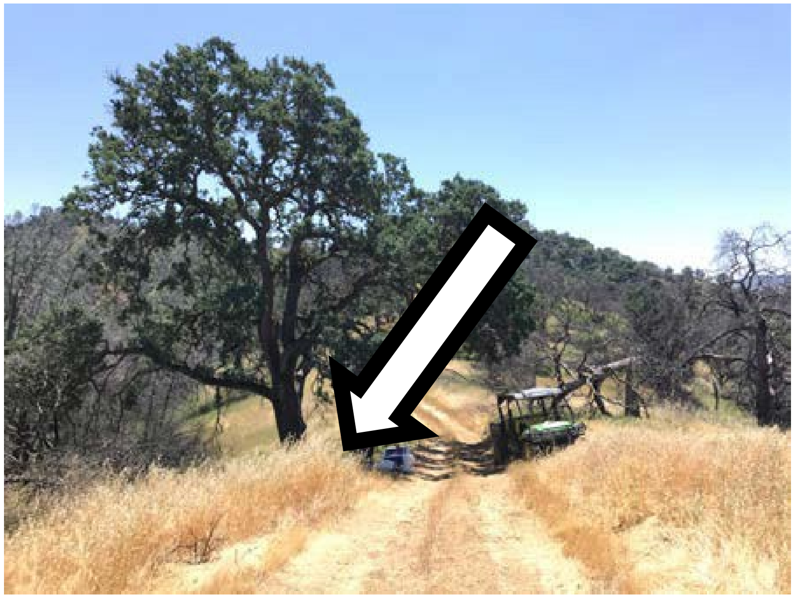
Tree 04
