## Supplementary Table 2 for "Functional composition of subsoil microbial communities changes with oak mortality"

| DO_ID | F1 | F2 |
|---|---|---|
| DO_T02_0-7.5 | 1.449526 | 0.136505 |
| DO_T02_7.5-15 | -0.36559 | -1.66525 |
| DO_T02_15-22.5 | -0.30893 | -0.82649 |
| DO_T02_22.5-30 | -0.01524 | -0.04955 |
| DO_T02_30-37.5 | -0.48072 | -0.25463 |
| DO_T02_37.5-45 | -1.15168 | -0.64303 |
| DO_T03_0-7.5 | 0.954656 | -1.61303 |
| DO_T03_7.5-15 | -0.15472 | -1.02254 |
| DO_T03_15-22.5 | -0.55477 | -1.4706 |
| DO_T03_22.5-30 | -0.29797 | -0.37062 |
| DO_T03_30-37.5 | -0.47929 | -0.01404 |
| DO_T03_37.5-45 | -0.98852 | 0.109022 |
| DO_T04_0-7.5 | 2.830337 | -0.00799 |
| DO_T04_7.5-15 | -0.09213 | -1.03653 |
| DO_T04_15-22.5 | -1.17961 | -1.46229 |
| DO_T04_22.5-30 | -0.82117 | 0.273441 |
| DO_T04_30-37.5 | -0.74802 | 2.038212 |
| DO_T04_37.5-45 | -0.89983 | 1.63813 |
| DO_T05_0-7.5 | 1.24046 | -0.27291 |
| DO_T05_7.5-15 | -0.21119 | -1.24824 |
| DO_T05_15-22.5 | -0.21862 | -0.07938 |
| DO_T05_22.5-30 | -1.26772 | -1.08365 |
| DO_T05_30-37.5 | -0.67698 | 0.239451 |
| DO_T05_37.5-45 | -0.98111 | 0.438462 |
| DO_T06_0-7.5 | 1.982837 | -0.31318 |
| DO_T06_7.5-15 | 0.221824 | -0.18987 |
| DO_T06_15-22.5 | 1.156625 | 2.857356 |
| DO_T06_22.5-30 | -0.07551 | 1.154385 |
| DO_T06_30-37.5 | -0.85241 | 0.19327 |
| DO_T06_37.5-45 | 0.325158 | 2.79234 |
| DO_T07_0-7.5 | 2.915412 | 0.269512 |
| DO_T07_7.5-15 | 0.978432 | 0.363179 |
| DO_T07_15-22.5 | -0.15115 | 0.490292 |
| DO_T07_22.5-30 | 0.020668 | 0.295406 |
| DO_T07_30-37.5 | -0.58556 | 0.30993 |
| DO_T07_37.5-45 | -0.51749 | 0.024935 |

Supplementary Table 2. Results of exploratory factor analysis of edaphic variables using fa() in R package psych. Based on a graph of factor correlations (Factor_Analysis_Graph_2factors), we estimate that F1 corresponds broadly to soil depth, while F2 corresponds to factors related to increased soil development (weathering).
